# Verification of SARS-CoV-2-Encoded small RNAs and contribution to Infection-Associated lung inflammation

**DOI:** 10.1101/2021.05.16.444324

**Authors:** Zhang Cheng, Liu Cheng, Jiang Lin, Cui Lunbiao, Li Chunyu, Song Guoxin, Xu Rui, Geng Xiangnan, Luan Changxing, Chen Yan, Zhu Baoli, Zhu Wei

**Affiliations:** Women & Children Central Laboratory, the First Affiliated Hospital of Nanjing Medical University, Nanjing Jiangsu 210036, China; Department of Gastroenterology, the First Affiliated Hospital of Nanjing Medical University, Nanjing Jiangsu 210029, China; Department of Endocrinology, the First Affiliated Hospital of Nanjing Medical University, Nanjing Jiangsu 210029, China; NHC Key Laboratory of Enteric Pathogen Microbiology, Jiangsu Provincial Center for Disease Control and Prevention, Nanjing Jiangsu 210009, China; Women & Children Intensive Care Unit, the First Affiliated Hospital of Nanjing Medical University, Nanjing Jiangsu 210036, China; Department of Pathology, the First Affiliated Hospital of Nanjing Medical University, Nanjing Jiangsu 210029, China; Emergency Medical Center, the First Affiliated Hospital of Nanjing Medical University, Nanjing Jiangsu 210029, China; Department of Clinical Engineering, the First Affiliated Hospital of Nanjing Medical University, Nanjing Jiangsu 210029, China; Department of Forensic Medicine, Nanjing Medical University, Nanjing Jiangsu 210029, China; Outpatient & Emergency Management Department, the First Affiliated Hospital of Nanjing Medical University, Nanjing, Jiangsu 210029, China; Jiangsu Provincial Center for Disease Control and Prevention, Nanjing Jiangsu 210009, China; Department of Oncology, the First Affiliated Hospital of Nanjing Medical University, Nanjing Jiangsu 210029, China

**Author notes:** **Corresponding Authors**: Zhu Wei, The First Affiliated Hospital of Nanjing Medical University, 300 Guangzhou Road, Nanjing, Jiangsu 210029, China.; Zhu Baoli, Jiangsu Provincial Center for Disease Control and Prevention, Nanjing Jiangsu 210009, China.; Chen Yan, The First Affiliated Hospital of Nanjing Medical University, 300 Guangzhou Road, Nanjing, Jiangsu 210029, China. Contributed equally (Zhang Cheng, Liu Cheng, Jiang Lin).

**Keywords:** SARS-CoV-2, COVID-19, small viral RNAs, inflammation

## Abstract

Severe acute respiratory syndrome coronavirus 2 (SARS-CoV-2) is the virus that causes coronavirus disease 2019 (COVID-19), the respiratory illness responsible for the COVID-19 pandemic. SARS-CoV-2 is a positive-stranded RNA virus belongs to *Coronaviridae* family. The viral genome of SARS-CoV-2 contains around 29.8 kilobase with a 5′-cap structure and 3′-poly-A tail, and shows 79.2% nucleotide identity with human SARS-CoV-1, which caused the 2002-2004 SARS outbreak. As the successor to SARS-CoV-1, SARS-CoV-2 now has circulated across the globe. There is a growing understanding of SARS-CoV-2 in virology, epidemiology, and clinical management strategies. In this study, we verified the existence of two 18-22 nt small viral RNAs (svRNAs) derived from the same precursor in human specimens infected with SARS-CoV-2, including nasopharyngeal swabs and formalin-fixed paraffin-embedded (FFPE) explanted lungs from lung transplantation of COVID-19 patients. We then simulated and confirmed the formation of these two SARS-CoV-2-Encoded small RNAs in human lung epithelial cells. And the potential pro-inflammatory effects of the splicing and maturation process of these two svRNAs in human lung epithelial cells were also explored. By screening cytokine storm genes and the characteristic expression profiling of COVID-19 in the explanted lung tissues and the svRNAs precursor transfected human lung epithelial cells, we found that the maturation of these two small viral RNAs contributed significantly to the infection associated lung inflammation, mainly via the activation of the CXCL8, CXCL11 and type I interferon signaling pathway. Taken together, we discovered two SARS-CoV-2-Encoded small RNAs and investigated the pro-inflammatory effects during their maturation in human lung epithelial cells, which might provide new insight into the pathogenesis and possible treatment options for COVID-19.

## 1. Introduction

Severe acute respiratory syndrome coronavirus 2 (SARS-CoV-2) is the virus that causes coronavirus disease 2019 (COVID-19), the respiratory illness responsible for the COVID-19 pandemic ^[1]^. SARS-CoV-2 belongs to the *Coronaviridae* family, a group of viruses co-infecting humans and other vertebrate animals. Coronaviruses (CoVs) infections affect the respiratory, gastrointestinal, liver, and central nervous systems of humans, livestock, birds, bats, mice, and many other wild animals. For instance, severe acute respiratory syndrome (SARS) in 2002 and the Middle East respiratory syndrome (MERS) in 2012 were both coronaviruses that transmitted from animals to humans. As a new evolutionary branch within the CoVs, SARS-CoV-2 contains around 29.8 kilobase with a 5′-cap structure and 3′-poly-A tail and shows 79.2% nucleotide identity with human SARS-CoV-1, which caused SARS outbreak ^[2]^. As the successor to SARS-CoV-1, SARS-CoV-2 shares a highly similar gene sequence and behavior pattern with SARS-CoV-1. Both of them are homologous RNA viruses and can transmit from person to person, causing a variety of diseases including pneumonia, hepatitis, encephalomyelitis, nephritis, enteritis and other illness. Despite the above similarities, there are many differences between SARS-CoV-1 and SARS-CoV-2, such as infectivity, latency, virulence difference, exposure dose, the rate of mutation, propagation velocity, super transmission rate, case fatality rate, susceptible population, stability, immunogenicity, treatment strategy, etc ^[3]^.

Up to the present, SARS-CoV-2 is the seventh human-transmitted coronavirus. SARS-CoV-1, MERS-CoV and SARS-CoV-2 can cause severe disease, whereas HKU1, NL63, OC43 and 229E are associated with mild symptoms ^[4]^. Data shows that the majority of patients with COVID-19 are asymptomatic or have mild symptoms, approximately 16 to 19% of patients develop acute respiratory failure, and 0.4 to 11.1% die from the disease ^[5]^. The primary targets of SARS-CoV-2 are the type-II alveolar epithelial cells and airway-epithelial cells, which highly express the angiotensin converting enzyme-2 (ACE2) receptor on their surface. After spike (S) glycoprotein binding to its receptor (ACE2) on the epithelial cells of the respiratory tracts, facilitated by cellular proteases furin and TMPRSS2, SARS-CoV-2 enters and quickly replicates inside the cells and kick-start the plethora of signaling cascade, from activating the pro-inflammatory effects to antiviral response leading to cytokine storm ^[6]^. The virus rapidly disseminates through peripheral blood to other organs like, heart, kidney, liver, spleen, etc. Immune responses to viral infection have evolved to clear the pathogen, and differences in these responses amongst patients probably affects clinical manifestations and outcomes ^[7, 8]^. Although there has been significant progress in understanding the factors involved with SARS-CoV-2 cellular infectivity, the relationship of SARS-CoV-2 lung infection and severity of pulmonary disease manifestations is not fully understood. Current studies revealed that dysregulated cellular immune responses and cytokine storm, e.g., aggregation of inflammatory monocyte-derived macrophages, lower CD8+ T cell infiltration and higher expression levels of cytokines (interleukins, interferons, tumor necrosis factors, colony stimulation factors, etc) and correlated receptors and proteins, might play important roles in the development of severe disease ^[9–11]^.

There are four major structural proteins encoded by SARS-CoV-2 genome: spike (S), nucleocapsid (N), membrane (M) and envelope (E), which are essential components of the virus particles ^[12, 13]^. It has been known that the E protein of SARS-CoV-1, being a virulence factor, participates in viral morphogenesis and contributes to the exacerbated inflammatory response associated with acute respiratory distress syndrome (ARDS) ^[14–16]^. In addition to protein components, a recent study has found that SARS-CoV-1 genome can also encode non-coding RNAs (ncRNAs) in the form of 18-22 nt small viral RNAs (svRNAs), which was independent of RNase III, cell type, and host species, but relying on the extent of viral replication ^[17]^. Although it is widely accepted that svRNAs can be derived from DNA viruses and RNA viruses with a nuclear stage, such as retroviruses, via the canonical small RNAs mature mechanism, growing evidence has supported that svRNAs could also be generated from cytoplasmic RNA viruses, which are relevant in the viral replication and pathogenicity ^[17–19]^. Exactly, as shown for SARS-CoV-1, inhibition of one of the svRNAs named svRNA-N, reduced in vivo lung pathology and pro-inflammatory cytokine expression were observed ^[17]^. The pathogenicity of SARS-CoV-2 is notably less than SARS-CoV-1 and MERS-CoV, but its high transmissibility led to the pandemic, which resulted in the global lock-down and affected the global health scenario adversely. In this study, we verified the existence of two SARS-CoV-2-Encoded small RNAs and investigated the pro-inflammatory effects during their maturation in human lung epithelial cells, which might provide new insight into the pathogenesis and possible treatment options for COVID-19.

## 2. Materials and Methods

### 2.1 Sample

The nucleic acid samples from pharyngeal swabs of 5 patients with COVID-19 were obtained from Jiangsu Provincial Center for Disease Control and Prevention from March 5 to April 7, 2020. The formalin-fixed paraffin-embedded (FFPE) explanted lung tissues from lung transplantation of two COVID-19 patients were provided by Department of forensic medicine, School of basic medicine, Nanjing Medical University, from August 3 to November 6, 2020. Three formalin-fixed paraffin-embedded (FFPE) non-infected lung tissues served as controls were from Department of pathology, the First Affiliated Hospital of Nanjing Medical University, from May 3 to June 1, 2020.

### 2.2 Basic local alignment search tool (BLAST) analysis

Previous study has identified 10 most abundant small viral RNAs (svRNAs) via deep sequencing in SARS-CoV-1 infected lungs derived from a mouse model ^[17]^. Among these 10 most abundant small viral RNAs (svRNAs), finally 3 svRNAs were characterized and only one of them, whose biogenesis and existence remarkably contributed to lung pathology and expression of pro-inflammatory cytokines in vivo. Based on the findings of the inflammatory SARS-CoV-1-Encoded small RNAs and the genome homology between SARS-CoV-1 and SARS-CoV-2, we hypothesized that small viral RNAs (svRNAs) encoded by SARS-CoV-2 not only exist but also share a highly similar gene sequence with those encoded by SARS-CoV-1. Hence, we compared the above 10 SARS-CoV-1-Encoded svRNAs with the SARS-CoV-2 genome via basic local alignment search tool (BLAST) analysis and proposed six potential small viral RNAs (svRNAs) encoded by SARS-CoV-2. These 6 potential svRNAs derived from the SARS-CoV-2 genome showed significant similarity with those encoded by SARS-CoV-1 in both nucleic acid sequences and virus genome positions, which were served as candidate SARS-CoV-2-Encoded svRNAs for further verification (supplementary Table S1).

### 2.3 Poly(A) RT-PCR combined with Pyrosequencing

The nucleic acid samples from pharyngeal swabs of 5 patients with COVID-19 were used for the screening and identification of the above six potential small viral RNAs (svRNAs) encoded by SARS-CoV-2, which were proposed by BLAST analysis. The RNA underwent Poly(A) tailing, reverse transcription and PCR using the miDETECT A Track™ miRNA RT-PCR Kit (Ribobio) containing the commercial miDETECT A Track™ Uni-RT primer and Uni-Reverse primer, while the forward primers for the above six potential svRNAs encoded by SARS-CoV-2 were synthesized by Tsingke Biotechnology and listed in supplementary Table S2. The PCR reactions were run on the qTOWER³ 84 (Analytik Jena) at 95□°C for 20□s, followed by 40□cycles of 10 s at 95□°C, 20□s at 60□°C. Finally the PCR products were purified from agarose gels and pyrosequenced by Tsingke Biotechnology to verify the existence of the above six potential svRNAs derived from SARS-CoV-2 genome.

### 2.4 Stem-loop RT-PCR combined with Pyrosequencing

Total RNA of formalin-fixed paraffin-embedded (FFPE) explanted lungs from lung transplantation of two COVID-19 patients was extracted through RNAprep Pure FFPE Kit (TIANGEN), and used for the further verification of the two identified SARS-CoV-2-Encoded svRNAs. To simulate and confirm the formation of the two SARS-CoV-2-Encoded small RNAs in human lung epithelial cells, total RNA of human lung epithelial cells transfected with the precursor of the two svRNAs was isolated, via TRIzol reagent (Invitrogen/Life Technologies, Carlsbad, California) following the manufacturer’s protocol. After the preparation of the RNA, SYBR Green based stem-loop RT-PCR detection was then used to confirm the specific 3’ splice sites during the mature of the two identified SARS-CoV-2-Encoded svRNAs. The stem-loop RT primers and PCR primers for svRNAs were synthesized by Tsingke Biotechnology and listed in supplementary Table S3. The PCR reactions were run on the LineGene 9600 (Bioer Technology) at 95□°C for 20□s, followed by 40□cycles of 10□s at 95□°C, 20□s at 60□°C. Finally the PCR products were purified from agarose gels and pyrosequenced by Tsingke Biotechnology to check the existence of the two verified svRNAs derived from SARS-CoV-2 genome.

### 2.5 In vitro RNA synthesis and purification

The precursor of the two verified SARS-CoV-2-Encoded svRNAs and negative control endogenous miRNA precursors were synthesized by in vitro transcription using T7 RNA polymerase with synthetic DNA templates obtained from Tsingke Biotechnology. The sequences of the control miRNA precursors were obtained from miRbase (http://www.mirbase.org). The sequences of sense and antisense single-stranded DNA (ssDNA) were listed in supplementary Table S4. Two ssDNA containing T7 promoter were mixed in a 1:1 ratio, denatured at 95 °C for 10 min and gradually cooled down at room temperature to form the double-stranded DNA (dsDNA) template. Transcription reactions contained 200 mM Tris-HCl (pH 7.9), 30 mM MgCl2, 50 mM DTT, 50 mM NaCl, 10 mM spermidine, 5mM of each NTP, 20 U/μL T7 RNA Polymerase and 10 μM linearized double-stranded DNA (dsDNA) template. Reactions were incubated at 37 °C for 8 hours, added 1 μL (2 U) of DNase I then incubated at 37 °C for 15 min. Phenol: chloroform extraction followed by ethanol precipitation was carried out for RNA purification. The 260/280 and 260/230 ratios of absorbance values were used to assess the purity of RNA using a Nanodrop ND-1000 spectrophotometer (Thermo Fisher Scientific). The svRNA maturing from the 5’ end of the precursor (svRNA-5p) and that from the 3’ end (svRNA-3p) were also synthesized by Tsingke Biotechnology.

### 2.6 Cell culture and RNA transfection

16HBE (human bronchial epithelial cell line) cells were grown in Dulbecco’s modified Eagle’s medium (DMEM, GIBCO) with D-glucose, L-glutamine, sodium pyruvate, and 10% fetal bovine serum with Pennicillin/Streptomycin in an atmosphere containing 5% CO2 at 37°C. Before transfection, cells were seeded in 12-well plates. Precursors of short RNAs were transfected into 16HBE cells by Lipofectamine 2000 reagent (Invitrogen) according to the manufacturer’s instructions. Cells were then harvested 48, 96 hours post transfection for further RNA extraction and detection, respectively.

### 2.7 Characteristic expression profiling analysis of COVID-19

Gene expression profile datasets were searched from Gene Expression Omnibus (GEO) database(http://www.ncbi.nlm.nih.gov/geo/) of National Centre for Biotechnology Information (NCBI) (https://www.ncbi.nlm.nih.gov/) with the keyword “SARS-CoV-2 OR COVID-19”. A total of 9 datasets including 5 from human lung tissues, 3 from nasopharyngeal swabs, and 1 from tracheal aspirates were collected for further analysis.

The GEO accession IDs, sample characteristics and experiment types of the above 9 datasets were summerized in supplementary Table S5.

The “DESeq” R packages was used to analyze expression profiling data derived from high throughput sequencing. Significant differentially expressed genes (DEGs) between COVID-19 and healthy control samples were firstly identified with the criteria of P-value < 0.01 and fold change > 2.0 or < 0.5. DEGs generated from each dataset were then compared and intersected to obtain a comprehensive DEGs list that pointed to be the characteristic expression profiling of COVID-19.

### 2.8 Reverse transcription quantitative PCR (RT-qPCR) assay

Reverse transcription quantitative PCR (RT-qPCR) assay for 23 cytokine genes and the above characteristic expression profiling of COVID-19 in different sample sources was used to investigate the pro-inflammatory effects of SARS-CoV-2 and the contribution to infection-associated lung inflammation caused by the maturation of the two SARS-CoV-2-Encoded svRNAs.

The 23 cytokine genes covered most of the members of the cytokine storm which were rapidly produced in human body fluid after infection with microorganisms, including chemokines (CXCL1, CXCL2, CXCL3, CXCL5, CXCL6, CXCL8, CXCL9, CXCL10, CXCL11, CXCL16), granulocyte colony-stimulating factor (G-CSF), granulocyte-macrophage colony stimulating factor (GM-CSF), interferons (IFNA1, IFNB1, IFNG, IFNL1), interleukins (IL1A, IL1B, IL2, IL27, IL6), tumor-necrosis factors (TNFA) and transforming-growth factor beta (TGFB).

Reverse transcription and quantitative PCR reactions were performed using PrimeScript RT reagent Kit (Takara) and SYBR Premix Ex Taq II (Takara) following the manufacturer’s protocols, respectively. The PCR reactions were run on the qTOWER³ 84 (Analytik Jena) at 95□°C for 20□s, followed by 40□cycles of 10□s at 95□°C, 20□s at 60□°C. The sequences of PCR primers were listed in the supplementary Table S6 and supplementary Table S7. GAPDH and 18sRNA were considered as reference genes for normalization, and the 2^−^△△Ct method was used to analyze the relative expression of target genes.

## 3 Results

### 3.1 The discovery of the two svRNAs encoded by SARS-CoV-2

As mentioned above, six potential small viral RNAs (svRNAs) encoded by SARS-CoV-2 were proposed, because of the significant similarity with those encoded by SARS-CoV-1 in both nucleic acid sequences and virus genome positions. The Poly(A) RT-PCR combined with Pyrosequencing in 5 nucleic acid samples from pharyngeal swabs of COVID-19 patients obtained from Jiangsu Provincial Center for Disease Control and Prevention showed that two svRNAs were really detected. Among the six potential svRNAs, the two svRNAs probably maturing from the same precursor named as svRNA-5p and svRNA-3p were firstly detected via Poly(A) RT-PCR with the amplification and melting curves (supplementary Figure S1A). Then the pyrosequencing of the PCR products showed that the two svRNAs really existed in the nucleic acid samples from pharyngeal swabs of COVID-19 patients, especially the svRNA-5p, which was found in all the 5 nucleic acid samples (Figure 1A). Thus, we believed that two svRNAs probably maturing from the same precursor encoded by SARS-CoV-2 existed.

**Figure 1:**
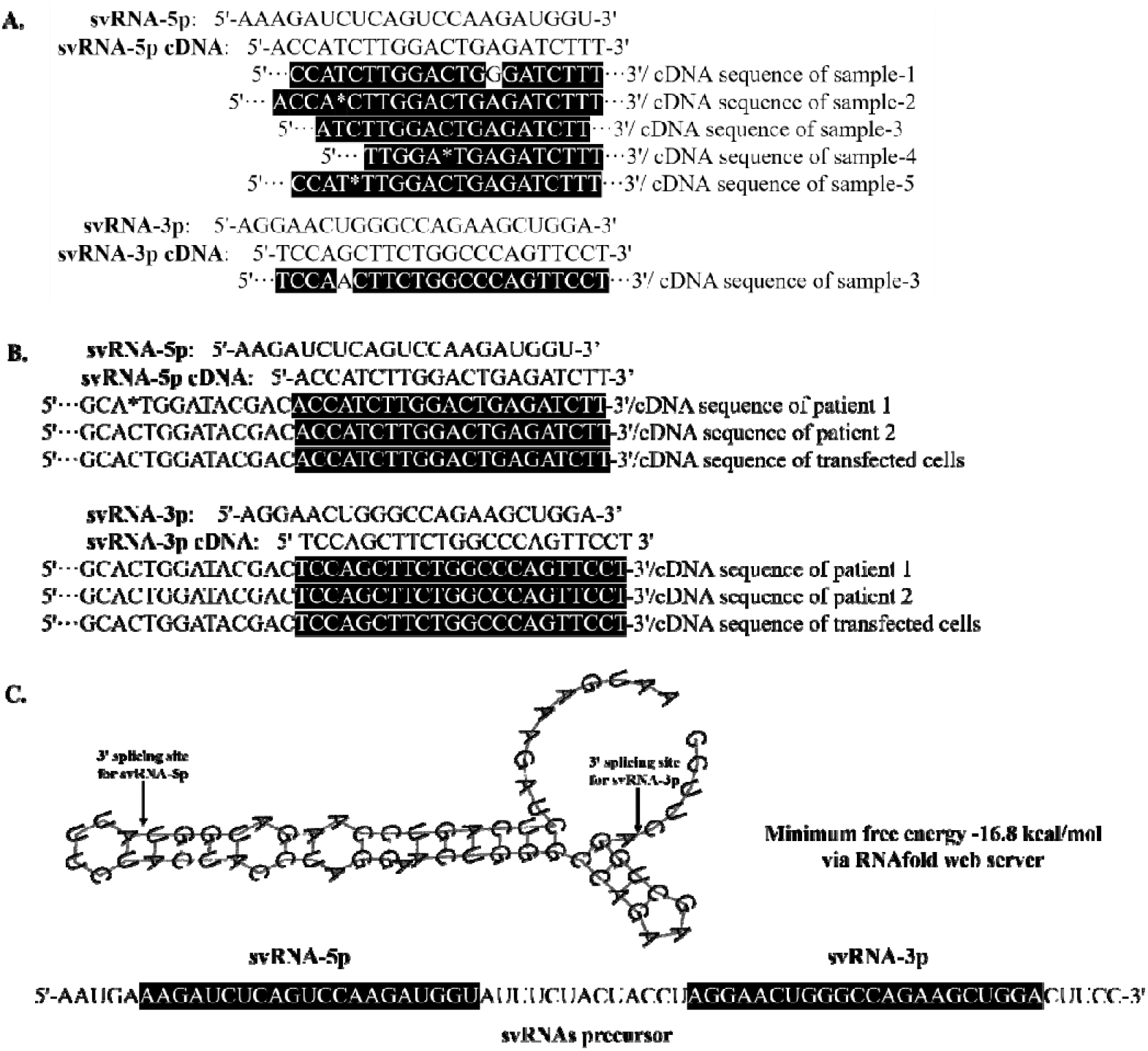
Sequences of two svRNAs encoded by SARS-CoV-2. (A) The pyrosequencing of the poly(A) RT-PCR products showed that two svRNAs encoded by SARS-CoV-2 (svRNA-5p and svRNA-3p), existed in the 5 nucleic acid samples from pharyngeal swabs of COVID-19 patients, especially the svRNA-5p, which was found in all the 5 nucleic acid samples. (B) The pyrosequencing of stem-loop RT-PCR products in FFPE explanted lungs of two COVID-19 patients and 16HBE (human lung epithelial cell line) cells transfected with the precursor of the two svRNAs further confirmed the existence of the two svRNAs encoded by SARS-CoV-2. (C) The secondary structure of svRNAs precursor with the length of 66 Bp via RNAfold web server based on minimum free energy.

### 3.2 The verification of the two svRNAs encoded by SARS-CoV-2

To further confirm the existence of the two svRNAs encoded by SARS-CoV-2, RNA of formalin-fixed paraffin-embedded (FFPE) explanted lungs from lung transplantation of two COVID-19 patients and three FFPE non-infected lung tissues which served as controls were extracted and used. SYBR Green based stem-loop RT-PCR detection firstly confirmed the specific 3’ splice sites during the mature of the two identified SARS-CoV-2-Encoded svRNAs in the lung of two COVID-19 patients. The corresponding amplification and melting curves were shown in the supplementary Figure S1B. There were no amplification and melting curves for all the three non-infected lung tissues (data not shown). Then the PCR products were purified from agarose gels (shown in supplementary Figure S1C) and pyrosequenced. The sequence of the PCR products verified that the two svRNAs encoded by SARS-CoV-2, probably maturing from the same precursor named as svRNA-5p and svRNA-3p, really existed (Figure 1B).

Moreover, to simulate and clarify the formation of the two SARS-CoV-2-Encoded small RNAs in human lung epithelial cells, RNA of human lung epithelial cells transfected with the precursor of the two svRNAs or the control short RNA precursors were isolated and used. SYBR Green based stem-loop RT-PCR combined with pyrosequencing also proved that the two SARS-CoV-2-Encoded small RNAs could splice and mature from the same precursor in the human lung epithelial cells. The corresponding amplification and melting curves were shown in the supplementary Figure S1B. There were no amplification and melting curves for those human lung epithelial cells transfected with the control short RNA precursors (data not shown). The PCR products purified from agarose gels was shown in supplementary Figure S1C, while the sequence of the PCR products was shown in the Figure 1B.

Finally, we tried to explore the secondary structure of svRNAs precursor which generated the two SARS-CoV-2 encoded svRNAs (svRNA-5p and svRNA-3p) via RNAfold web server based on minimum free energy. The secondary structure of svRNAs precursor with the different legths (from 56 bp to 96 bp) showed that the splicing sites of the two SARS-CoV-2 encoded svRNAs (svRNA-5p and svRNA-3p) were always stable, no matter with the length of svRNAs precursor (supplementary Figure S2), which also suggested that the existence of the two SARS-CoV-2 encoded svRNAs. The potential sequence of svRNAs precursor with different length for RNAfold web server to predict the secondary structures were shown in supplementary Table S8. The typical secondary structure of the two svRNAs precursor with the length of 66 Bp was also shown in the Figure 1C.

### 3.3 The characteristic expression profiling of COVID-19

Just as described above, 9 GEO datasets derived from high throughput sequencing, which were closely related with SARS-CoV-2 infection were collected and analyzed by the “DESeq” R packages. The significant DEGs between COVID-19 and healthy control samples were firstly identified. DEGs generated from each dataset were then compared and intersected to obtain a comprehensive DEGs list that pointed to be the characteristic expression profiling of COVID-19. As shown in the circular-barplot (Figure 2A), 35 genes were uniformly up-regulated in all 9 GEO datasets, while there were no genes down expressed consistently in all the 9 GEO datasets. Thus, the 35 up-regulated genes upon SARS-CoV-2 infection in human respiratory tracts implied being the characteristic expression profiling of COVID-19.

**Figure 2:**
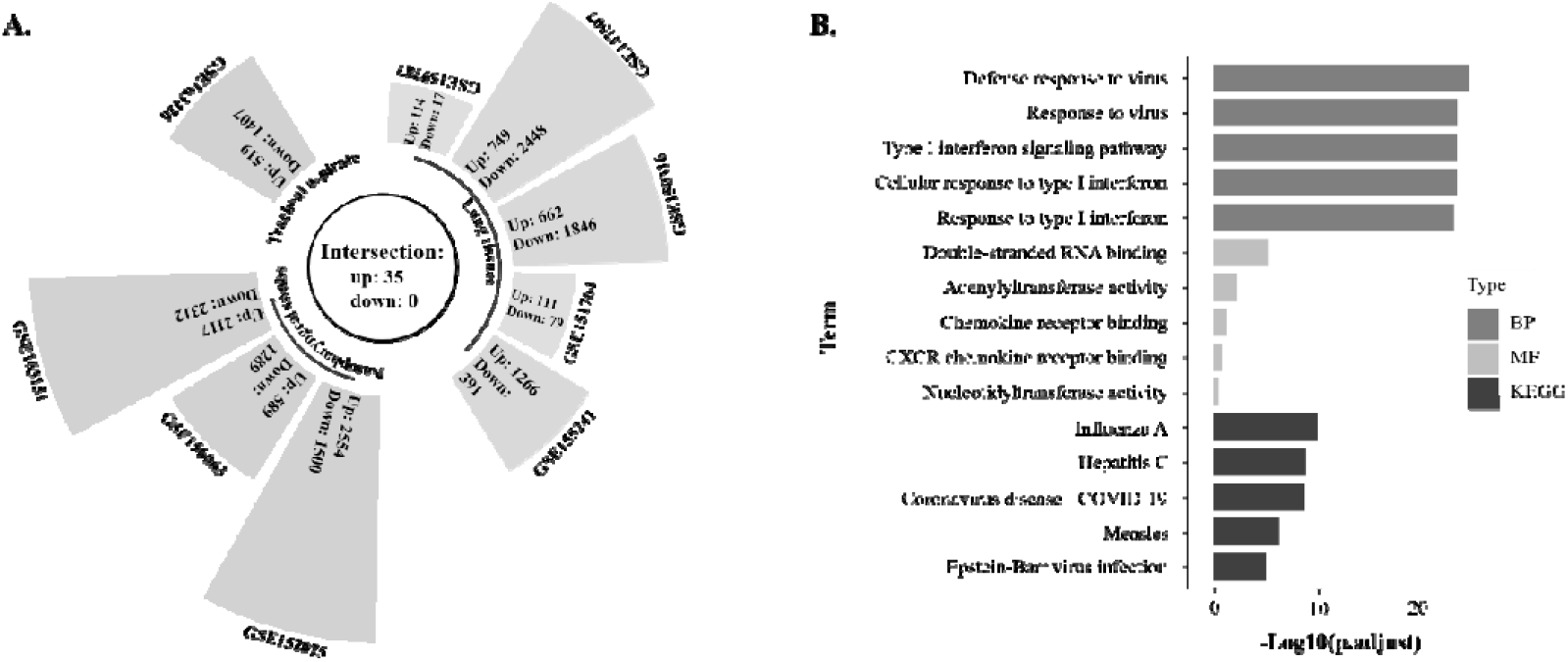
The characteristic expression profiling of COVID-19. (A) The circular-barplot showed 35 genes were uniformly up-regulated in all 9 GEO datasets including 5 from human lung tissues, 3 from nasopharyngeal swabs, and 1 from tracheal aspirates. For down-regulated genes, there was no concordance across these datasets. Thus, the 35 up-regulated genes upon SARS-CoV-2 infection in human respiratory tracts implied being the characteristic expression profiling of COVID-19. (B) Gene Ontology (GO) biological process (BP), molecular function (MF) and Kyoto Encyclopedia of Genes and Genomes (KEGG) functional enrichment analysis of the characteristic expression profiling of COVID-19, namely the 35 significantly up-regulated genes, showed that type I interferon biological process and chemokine molecular activation, ranked in the top 5 of the infection profiles filtered by adjust P-value, were obviously involved in the SARS-CoV-2 caused inflammation.

Gene Ontology (GO) biological process (BP), molecular function (MF) and Kyoto Encyclopedia of Genes and Genomes (KEGG) functional enrichment analysis of the characteristic expression profiling of COVID-19, namely the 35 significantly up-regulated genes, showed that type I interferon biological process and chemokine molecular activation, ranked in the top 5 of the infection profiles filtered by adjust P-value, were obviously involved in the SARS-CoV-2 caused inflammation (Figure 2B).

### 3.4 The pulmonary inflammation feature analysis of COVID-19

To systematically investigate the pulmonary inflammation feature of COVID-19 at transcriptional level, 23 cytokine genes covering most of the members of the cytokine storm which were rapidly produced in human body fluid after infection with microorganisms, and 35 significantly up-regulated genes, namely the characteristic expression profiling of COVID-19, were included for RT-qPCR assay analysis between the FFPE explanted lung tissues from lung transplantation of two COVID-19 patients and three FFPE non-infected lung tissues which served as controls. Among the 23 cytokine genes, 8 genes including chemokines (CXCL5, CXCL9, CXCL11), interferons (IFNG, IFNL1), G-CSF, GM-CSF and IL1A were over-expressed consistently in both two COVID-19 patients when compared with the controls, respectively (Figure 3A).

**Figure 3:**
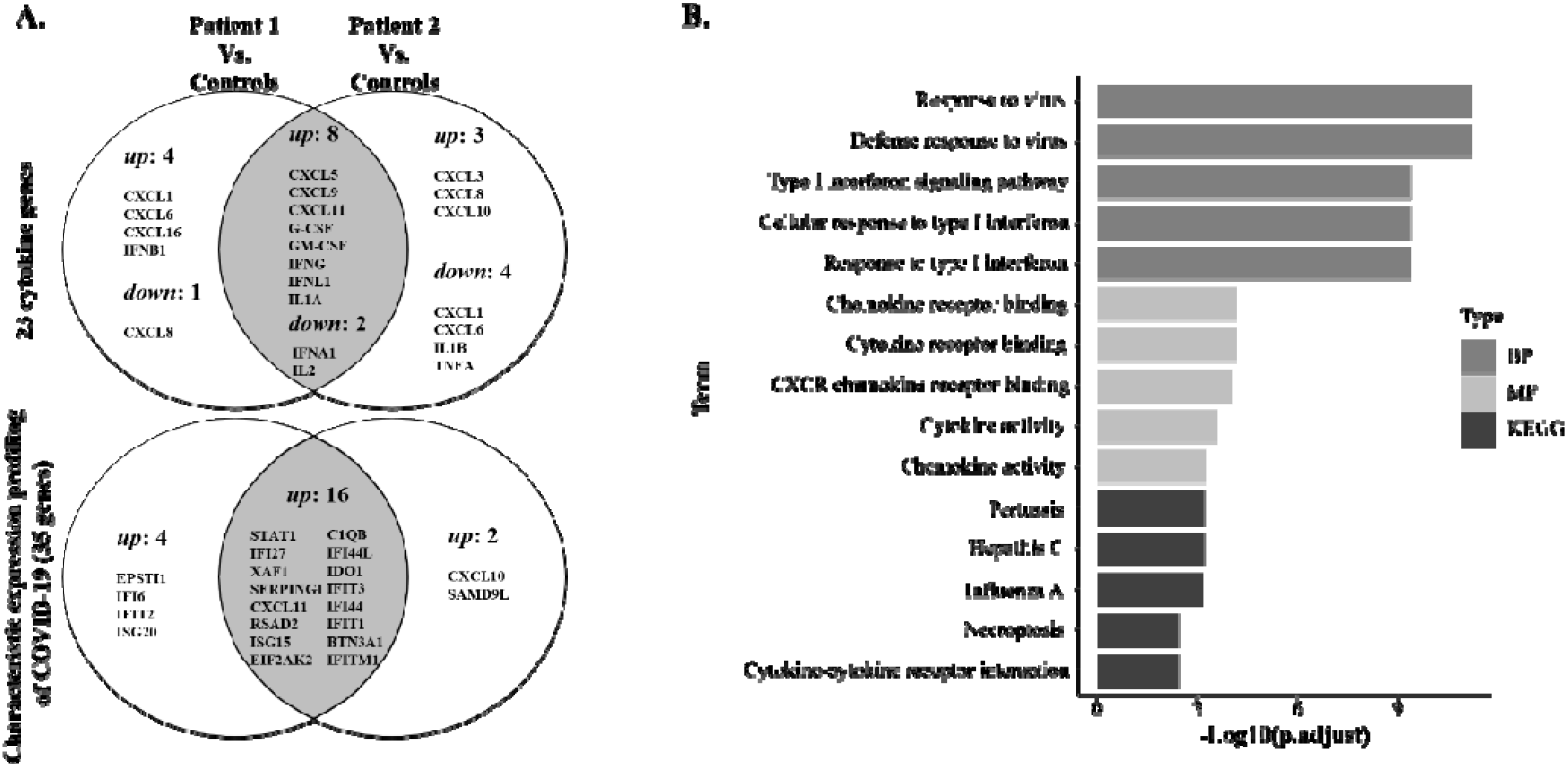
The pulmonary inflammation feature analysis of COVID-19. (A) RT-qPCR assay analysis of 23 cytokine genes and 35 genes, namely the characteristic expression profiling of COVID-19, between the FFPE explanted lung tissues from lung transplantation of two COVID-19 patients and three FFPE non-infected control lung tissues, showed that 8 cytokine genes including chemokines (CXCL5, CXCL9, CXCL11), interferons (IFNG, IFNL1), G-CSF, GM-CSF, IL1A and 16 genes from the characteristic expression profiling of COVID-19, were over-expressed consistently in both two COVID-19 patients when compared with the controls, respectively. (B) Gene Ontology (GO) biological process (BP), molecular function (MF) and Kyoto Encyclopedia of Genes and Genomes (KEGG) functional enrichment analysis of the these 23 commonly up-regulated genes in the lungs of COVID-19 patients showed that type I interferon biological process and chemokine molecular activation were ranked in the top 5 of the infection profiles filtered by adjust P-value. Hence, the type I interferon biological process and chemokine molecular activation at least in part consisted of the pulmonary inflammation feature of COVID-19.

As to the 35 genes of characteristic expression profiling of COVID-19, 16 genes were commonly up-regulated in both two COVID-19 patients (Figure 3A). All the differential expression analysis results of the 23 cytokine genes and the 35 genes of characteristic expression profiling of COVID-19 in the lung tissues between the two COVID-19 patients and the controls were shown in supplementary Table S9 and Table S10, respectively.

Gene Ontology (GO) biological process (BP), molecular function (MF) and Kyoto Encyclopedia of Genes and Genomes (KEGG) functional enrichment analysis of the these 23 commonly up-regulated genes in the lungs of COVID-19 patients also showed that type I interferon biological process and chemokine molecular activation were ranked in the top 5 of the infection profiles filtered by adjust P-value (Figure 3B). Hence, we believed that type I interferon biological process and chemokine molecular activation at least in part consisted of the pulmonary inflammation feature of COVID-19.

### 3.5 Exploration of the pro-inflammatory effects of SARS-CoV-2 encoded svRNAs

By screening 23 cytokine storm genes and the characteristic expression profiling of COVID-19, we found that type I interferon biological process and chemokine molecular activation were the underlying mechanism of the SARS-CoV-2 infection associated lung inflammation. To explore the pro-inflammatory effects of SARS-CoV-2 encoded svRNAs in human lung epithelial cells, the mature and the precursor svRNAs were transfected into cells and the RT-qPCR assay was then used to analysize the expression levels of those lung inflammation associated genes. As shown in Figure 4A, those transfected with the precursor of the svRNAs exhibited the most remarkable inflammatory reaction at transcriptional level, no matter compared with those transfected with mature svRNAs or those transfected with endogenous short RNA precursors. However, those transfected with the mature svRNAs had no distinguishing inflammatory reaction, suggesting that the pro-inflammatory effects of double-stranded RNA (dsRNA) was stronger than that of single-stranded RNA (ssRNA), which meant that the splicing and mature process of SARS-CoV-2 encoded svRNAs might closely related with SARS-CoV-2 infection associated lung inflammation. Exactly, Gene Ontology (GO) biological process (BP), molecular function (MF) and Kyoto Encyclopedia of Genes and Genomes (KEGG) functional enrichment analysis of the those up-regulated genes irritated by the svRNA precursor, compared with those transfected with endogenous short RNA precursors, still showed that type I interferon biological process and chemokine molecular activation were ranked in the top 5 of the infection profiles filtered by adjust P-value (Figure 4B).

**Figure 4:**
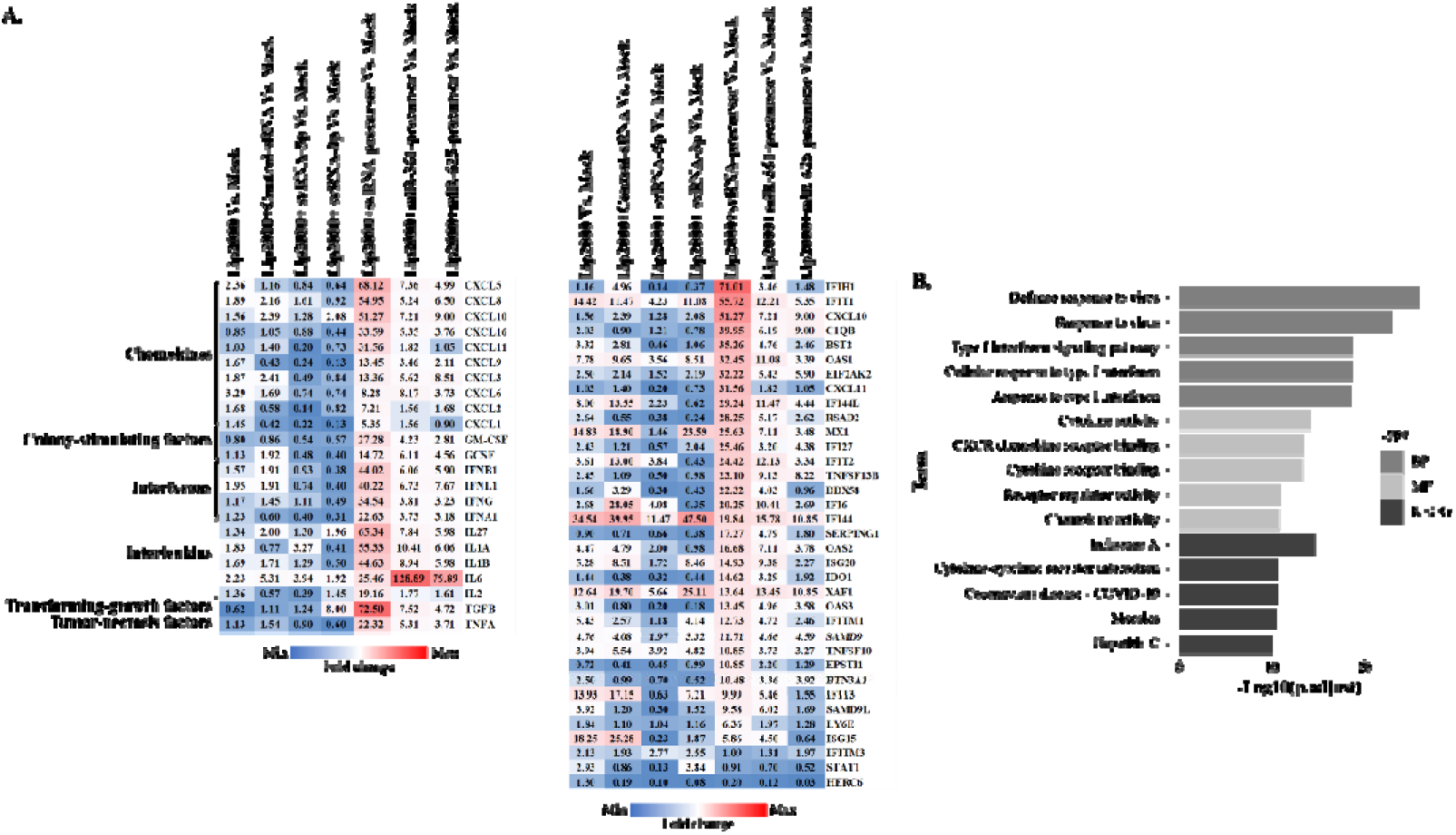
Exploration of the pro-inflammatory effects of SARS-CoV-2 encoded svRNAs. (A) 16HBE cells were separately transfected with 40 pmol control svRNA, control miR-361 precursor, control miR-625 precursor, svRNA-5p, svRNA-3p and svRNAs precursor. The heatmaps showed the gene expression in 16HBE cells 48 hours post-infection. The intensity of the color scheme was scaled to relative expression values (fold change). Those transfected with the precursor of the svRNAs exhibited the most remarkable inflammatory reaction at transcriptional level, no matter compared with those transfected with mature svRNAs or those transfected with endogenous short RNA precursors. (B) Gene Ontology (GO) biological process (BP), molecular function (MF) and Kyoto Encyclopedia of Genes and Genomes (KEGG) functional enrichment analysis of the those up-regulated genes irritated by the svRNA precursor, compared with those transfected with endogenous short RNA precursors, showed that type I interferon biological process and chemokine molecular activation were ranked in the top 5 of the infection profiles filtered by adjust P-value.

### 3.6 Contribution to inflammation of the svRNAs precursor

To further estimate the pro-inflammatory effects of the svRNA precursor, cells were transfected with exogenous and endogenous double-stranded short RNA precursors at different dosage levels and time courses, respectively. Within the 23 cytokine genes, chemokines (CXCL8, CXCL11, CXCL16) and interferons (IFNA1, IFNB1, IFNG, IFNL1) and 13 genes within the characteristic expression profiling of COVID-19, were over-expressed consistently at two dose levels, when compared with those transfected with endogenous short RNA precursors, respectively (Figure 5A). However, no obvious dose-dependence between svRNA precursor and inflammation reaction at transcriptional level was observed. All the RT-qPCR assay analysis on cells transfected with different dosage of short RNA precursors was shown in the Figure S3.

**Figure 5:**
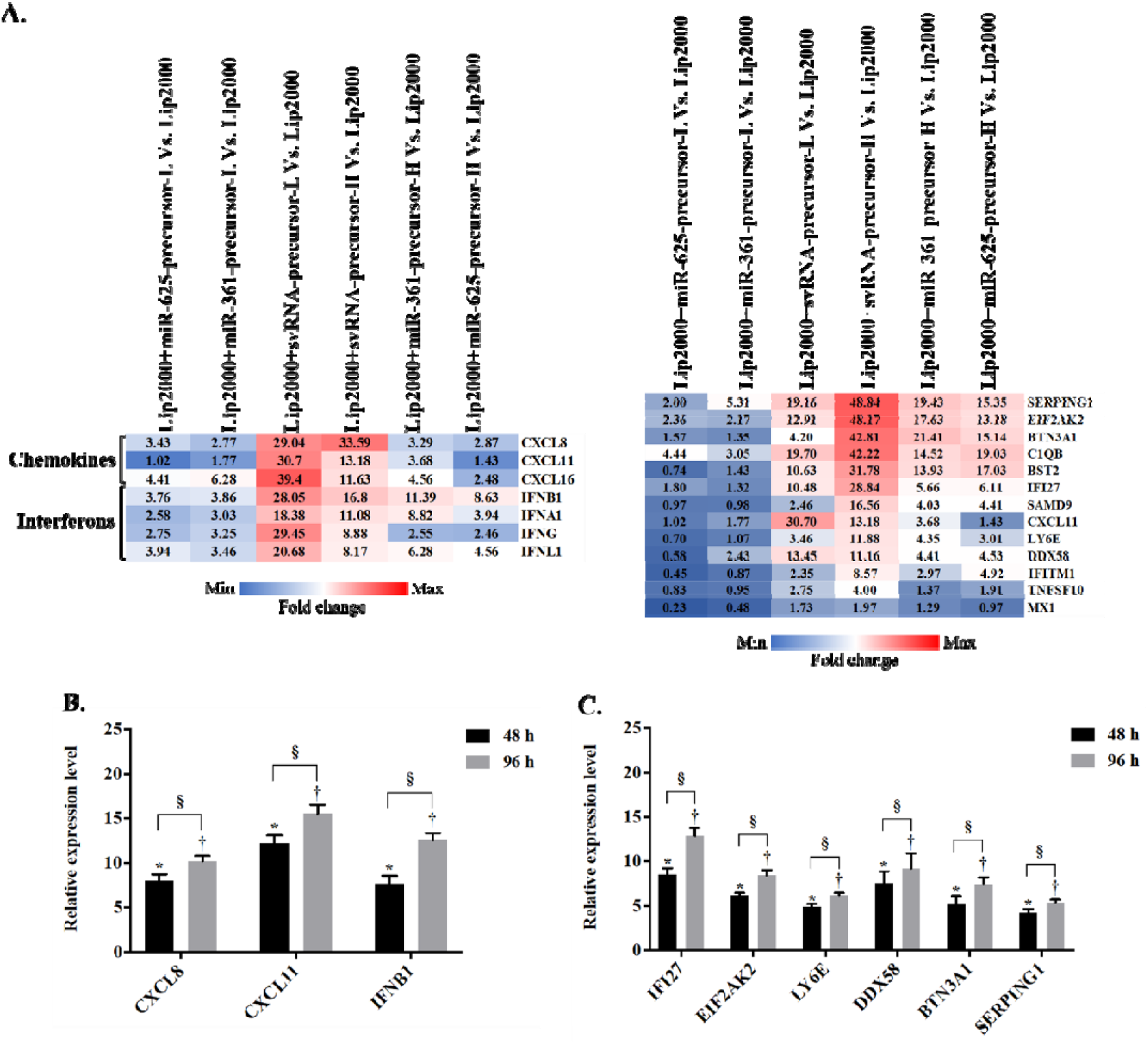
Contribution to inflammation of the svRNAs precursor. (A) 16HBE cells were transfected with control endogenous short RNA precursors and svRNAs precursor at the high (H) and low (L) dosage of 40 pmol and 80 pmol, respectively. The heatmaps showed the gene expression in 16HBE cells 48 hours post-infection. The intensity of the color scheme was scaled to relative expression values (fold change). Within the 23 cytokine genes, chemokines (CXCL8, CXCL11, CXCL16) and interferons (IFNA1, IFNB1, IFNG, IFNL1) and 13 genes within the characteristic expression profiling of COVID-19, were over-expressed consistently at two dose levels, when compared with those transfected with endogenous short RNA precursors, respectively. However, no obvious dose-dependence between svRNA precursor and inflammation reaction at transcriptional level was observed. (B&C) 16HBE cells were transfected with 40 pmol different short RNA precursors separately. Among those above 7 cytokine genes and 13 genes which were consistently up-regulated at different dosage levels, 3 cykokine genes of CXCL8, CXCL11, IFNB1 and 6 genes including IFI27, EIF2AK2, LY6E, DDX58, BTN3A1 and SERPING1, were both up-regulated in cells 48 hours and 96 hours post-transfection, compared with the control endogenous short RNA precursors, respectively. And the expression level of these genes were significantly higher in cells 96 hours than in those 48 hours post-transfection, which were shown to be time-dependent. Values are the mean ± standard deviation; n = 3; *: p < 0.05; †: p < 0.05; §: p < 0.05.

While as shown in Figure 5B and Figure 5C, among those above 7 cytokine genes and 13 genes within the characteristic expression profiling of COVID-19, which were consistently up-regulated at different dosage levels, 3 cykokine genes of CXCL8, CXCL11, IFNB1 and 6 genes including IFI27 (interferon-alpha (IFN-α)-inducible gene), EIF2AK2 (interferon-induced, eukaryotic translation initiation factor 2-alpha kinase 2), LY6E (lymphocyte antigen 6 family member E), DDX58 (antiviral innate immune response receptor RIG-I encoded gene), BTN3A1 (butyrophilin subfamily 3 member A1) and SERPING1 (plasma protease C1 inhibitor, serpin family G member 1), were both up-regulated in cells transfected with the svRNA precursor for 48 hours and 96 hours, respectively. And the expression level of these genes were significantly higher in cells 96 hours post-transfection than in those 48 hours post-transfection, which were shown to be time-dependent.

Taken all together, the maturation of these two SARS-CoV-2-Encoded small RNAs contributed significantly to the infection associated lung inflammation, mainly via the activation of the CXCL8, CXCL11 and type I interferon signaling pathway.

## 4 Discussion

This study aimed to discovery the small viral RNAs (svRNAs) encoded by SARS-CoV-2 genome. Based on the genome homology between SARS-CoV-1 and SARS-CoV-2, we compared the 10 most abundant SARS-CoV-1-encoded svRNAs identified in previous study with the SARS-CoV-2 genome and proposed six potential 18-22 nt svRNAs encoded by SARS-CoV-2. Among the six potential svRNAs, we verified the real existence of two svRNAs in human specimens infected with SARS-CoV-2, including nasopharyngeal swabs and formalin-fixed paraffin-embedded (FFPE) explanted lungs from lung transplantation of COVID-19 patients. Notably, these two svRNAs probably matured from the same precursor. We simulated and confirmed the formation of these two SARS-CoV-2-Encoded svRNAs in human lung epithelial cells. Moreover, the secondary structure of the svRNAs precursors with the different lengths (from 56 bp to 96 bp) predicted by RNAfold web server based on minimum free energy showed that the splicing sites of these two svRNAs were always stable, which also suggested the existence of the two SARS-CoV-2 encoded svRNAs. Then, we explored the pro-inflammatory effects of the SARS-CoV-2 encoded svRNAs and found that the maturation of them contributed significantly to the infection associated lung inflammation, mainly via the activation of the CXCL8, CXCL11 and type I interferon signaling pathway.

In the current study, we confirmed two SARS-CoV-2 encoded svRNAs out of six potential svRNA candidates. Not only for SARS-CoV-1 and SARS-CoV-2, plenty of RNA viruses, such as influenza virus, EV71, hepatitis A virus, HCV, Polio, Dengue, vesicular stomatitis, and West Nile viruses, have been found to generate svRNAs representing a low abundance but with relevant biological functions ^[17–20]^. It was worthy of note that these two SARS-CoV-2 encoded svRNAs might splice from the same precursor and located preferentially near the genome ends. This pattern was also observed in many other RNA viruses, suggesting that these svRNAs were not produced randomly or just degradation fragments from viral genome, but that they were generated specifically and played a role during infection.

Exactly, previous study had reported that the SARS-CoV-1 derived svRNAs were mainly generated independent of canonical cellular pathways, while were dependent on alternative mechanisms involving viral proteins or cellular factors induced during infection ^[21]^. However, the specific underlying mechanism was still unknown. In our study, by systematically estimating the pro-inflammatory effects of the SARS-CoV-2 encoded svRNAs in human lung epithelial cells, we found that the precursor of the svRNAs exhibited the most remarkable inflammatory reaction at transcriptional level while no distinguishing inflammatory reaction was provoked by the mature svRNAs. Moreover, the exogenous double-stranded RNA (dsRNA) precursor of SARS-CoV-2 encoded svRNAs contributed the similar pro-inflammatory effects with the SARS-CoV-2 virus did in the human lungs, mainly via the activation of the CXCL8, CXCL11 and type I interferon signaling pathway, which meant that the splicing and mature process of the SARS-CoV-2 encoded svRNAs might closely related with infection associated lung inflammation.

Specifically, 3 cytokine genes of CXCL8, CXCL11, IFNB1 and 6 genes from the characteristic expression profiling of COVID-19 including IFI27, EIF2AK2, LY6E, DDX58, BTN3A1 and SERPING1 were significantly induced at the transcriptional level by the SARS-CoV-2 encoded svRNAs precursor. Previous study had reported that high level of serum CXCL8 was positively correlated with the severity and mortality rate of COVID-19, while CXCL11 was the lung tissue-specific gene verified by single cell sequencing ^[22]^. As the member of the type I interferons, IFNB1 was expressed in most of epithelial cells, which had been reported to defense against SARS-CoV-2 infections in mild or moderate COVID-19 patients ^[23]^. It was also well known that the type I interferon signaling pathway could be activated by dsRNA ^[24, 25]^. Indeed, among the above 6 genes, 3 of them, FI27, EIF2AK2 and DDX58 were induced by type I interferon or involved in dsRNA process ^[26–28]^. In the rest of 3 genes, LY6E and BTN3A1 were associated with T-cell activation in the adaptive immune response, while SERPING1 was related to complement and coagulation cascades pathway ^[29–32]^. Especially for LY6E, recent studies had proved that LY6E restricted the entry of human coronaviruses mediated by the envelope spike proteins, including SARS-CoV-2 ^[31, 32]^.

Hence, this study here verified the existence of two SARS-CoV-2-Encoded svRNAs and explored the pro-inflammatory effects during their maturation in human lung epithelial cells. However, more research was needed on the mechanism of the biogenesis of SARS-CoV-2-Encoded svRNAs and the cell specificity of the generation of them. Moreover, the pro-inflammatory effects at different molecular levels during the maturation of SARS-CoV-2-Encoded svRNAs were also required. Unraveling these molecular mechanisms of SARS-CoV-2-Encoded svRNAs would be helpful in the design of antivirals to control the exacerbated host immune response induced by SARS-CoV-2 infection.

## Competing financial interests

The authors declare no competing financial interests.

## Acknowledgements

The authors would like to thank Professor of Cheng Feng for his support on providing the FFPE lung tissues of COVID-19 patients.

## Notes

### Competing Interest Statement

The authors have declared no competing interest.

## References

[1] Wang Y, Zhang L, Sang L, Ye F, Ruan S, Zhong B, Song T, Alshukairi AN, Chen R, Zhang Z, Gan M, Zhu A, Huang Y, Luo L, Mok CKP, Al Gethamy MM, Tan H, Li Z, Huang X, Li F, Sun J, Zhang Y, Wen L, Li Y, Chen Z, Zhuang Z, Zhuo J, Chen C, Kuang L, Wang J, Lv H, Jiang Y, Li M, Lin Y, Deng Y, Tang L, Liang J, Huang J, Perlman S, Zhong N, Zhao J, Malik Peiris JS, Li Y, Zhao J. Kinetics of viral load and antibody response in relation to COVID-19 severity. J Clin Invest. 2020 Oct 1;130(10):5235–5244.

[2] Chia WN, Tan CW, Foo R, Kang AEZ, Peng Y, Sivalingam V, Tiu C, Ong XM, Zhu F, Young BE, Chen MI, Tan YJ, Lye DC, Anderson DE, Wang LF. Serological differentiation between COVID-19 and SARS infections. Emerg Microbes Infect. 2020 Dec;9(1):1497–1505.

[3] Harrison AG, Lin T, Wang P. Mechanisms of SARS-CoV-2 Transmission and Pathogenesis. Trends Immunol. 2020 Dec;41(12):1100–1115.

[4] Li L, Zhang W, Hu Y, Tong X, Zheng S, Yang J, Kong Y, Ren L, Wei Q, Mei H, Hu C, Tao C, Yang R, Wang J, Yu Y, Guo Y, Wu X, Xu Z, Zeng L, Xiong N, Chen L, Wang J, Man N, Liu Y, Xu H, Deng E, Zhang X, Li C, Wang C, Su S, Zhang L, Wang J, Wu Y, Liu Z. Effect of Convalescent Plasma Therapy on Time to Clinical Improvement in Patients With Severe and Life-threatening COVID-19: A Randomized Clinical Trial. JAMA. 2020 Aug 4;324(5):460–470.

[5] Huang C, Wang Y, Li X, Ren L, Zhao J, Hu Y, Zhang L, Fan G, Xu J, Gu X, Cheng Z, Yu T, Xia J, Wei Y, Wu W, Xie X, Yin W, Li H, Liu M, Xiao Y, Gao H, Guo L, Xie J, Wang G, Jiang R, Gao Z, Jin Q, Wang J, Cao B. Clinical features of patients infected with 2019 novel coronavirus in Wuhan, China. Lancet. 2020 Feb 15;395(10223):497–506.

[6] Lother SA, Abassi M, Agostinis A, Bangdiwala AS, Cheng MP, Drobot G, Engen N, Hullsiek KH, Kelly LE, Lee TC, Lofgren SM, MacKenzie LJ, Marten N, McDonald EG, Okafor EC, Pastick KA, Pullen MF, Rajasingham R, Schwartz I, Skipper CP, Turgeon AF, Zarychanski R, Boulware DR. Post-exposure prophylaxis or pre-emptive therapy for severe acute respiratory syndrome coronavirus 2 (SARS-CoV-2): study protocol for a pragmatic randomized-controlled trial. Can J Anaesth. 2020 Sep;67(9):1201–1211.

[7] Secchi M, Bazzigaluppi E, Brigatti C, Marzinotto I, Tresoldi C, Rovere-Querini P, Poli A, Castagna A, Scarlatti G, Zangrillo A, Ciceri F, Piemonti L, Lampasona V. COVID-19 survival associates with the immunoglobulin response to the SARS-CoV-2 spike receptor binding domain. J Clin Invest. 2020 Dec 1;130(12):6366–6378.

[8] Carsetti R, Zaffina S, Piano Mortari E, Terreri S, Corrente F, Capponi C, Palomba P, Mirabella M, Cascioli S, Palange P, Cuccaro I, Milito C, Zumla A, Maeurer M, Camisa V, Vinci MR, Santoro A, Cimini E, Marchioni L, Nicastri E, Palmieri F, Agrati C, Ippolito G, Porzio O, Concato C, Onetti Muda A, Raponi M, Quintarelli C, Quinti I, Locatelli F. Different Innate and Adaptive Immune Responses to SARS-CoV-2 Infection of Asymptomatic, Mild, and Severe Cases. Front Immunol. 2020 Dec 16;11:610300.

[9] Sattler A, Angermair S, Stockmann H, Heim KM, Khadzhynov D, Treskatsch S, Halleck F, Kreis ME, Kotsch K. SARS-CoV-2-specific T cell responses and correlations with COVID-19 patient predisposition. J Clin Invest. 2020 Dec 1;130(12):6477–6489.

[10] Zhou Q, Chen V, Shannon CP, Wei XS, Xiang X, Wang X, Wang ZH, Tebbutt SJ, Kollmann TR, Fish EN. Interferon-α2b Treatment for COVID-19. Front Immunol. 2020 May 15;11:1061.

[11] Varchetta S, Mele D, Oliviero B, Mantovani S, Ludovisi S, Cerino A, Bruno R, Castelli A, Mosconi M, Vecchia M, Roda S, Sachs M, Klersy C, Mondelli MU. Unique immunological profile in patients with COVID-19. Cell Mol Immunol. 2021 Mar;18(3):604–612.

[12] Satarker S, Nampoothiri M. Structural Proteins in Severe Acute Respiratory Syndrome Coronavirus-2. Arch Med Res. 2020 Aug;51(6):482–491.

[13] Hatmal MM, Alshaer W, Al-Hatamleh MAI, Hatmal M, Smadi O, Taha MO, Oweida AJ, Boer JC, Mohamud R, Plebanski M. Comprehensive Structural and Molecular Comparison of Spike Proteins of SARS-CoV-2, SARS-CoV and MERS-CoV, and Their Interactions with ACE2. Cells. 2020 Dec 8;9(12):2638.

[14] Afewerky HK. Pathology and pathogenicity of severe acute respiratory syndrome coronavirus 2 (SARS-CoV-2). Exp Biol Med (Maywood). 2020 Sep;245(15):1299–1307.

[15] Hu B, Guo H, Zhou P, Shi ZL. Characteristics of SARS-CoV-2 and COVID-19. Nat Rev Microbiol. 2021 Mar;19(3):141–154.

[16] V’kovski P, Kratzel A, Steiner S, Stalder H, Thiel V. Coronavirus biology and replication: implications for SARS-CoV-2. Nat Rev Microbiol. 2021 Mar;19(3):155–170.

[17] Morales L, Oliveros JC, Fernandez-Delgado R, tenOever BR, Enjuanes L, Sola I. SARS-CoV-Encoded Small RNAs Contribute to Infection-Associated Lung Pathology. Cell Host Microbe. 2017 Mar 8;21(3):344–355.

[18] Te Velthuis AJ, Fodor E. Influenza virus RNA polymerase: insights into the mechanisms of viral RNA synthesis. Nat Rev Microbiol. 2016 Aug;14(8):479–93.

[19] Cross ST, Michalski D, Miller MR, Wilusz J. RNA regulatory processes in RNA virus biology. Interdiscip Rev RNA. 2019 Sep;10(5):e1536.

[20] Mandary MB, Masomian M, Poh CL. Impact of RNA Virus Evolution on Quasispecies Formation and Virulence. Int J Mol Sci. 2019 Sep 19;20(18):4657.

[21] Nascimento Junior JAC, Santos AM, Quintans-Júnior LJ, Walker CIB, Borges LP, Serafini MR. SARS, MERS and SARS-CoV-2 (COVID-19) treatment: a patent review. Expert Opin Ther Pat. 2020 Aug;30(8):567–579.

[22] Espíndola OM, Gomes YCP, Brandão CO, Torres RC, Siqueira M, Soares CN, Lima MASD, Leite ACCB, Venturotti CO, Carvalho AJC, Torezani G, Araujo AQC, Silva MTT. Inflammatory Cytokine Patterns Associated with Neurological Diseases in Coronavirus Disease 2019. Ann Neurol. 2021 May;89(5):1041–1045.

[23] Park A, Iwasaki A. Type I and Type III Interferons - Induction, Signaling, Evasion, and Application to Combat COVID-19. Cell Host Microbe. 2020 Jun 10;27(6):870–878.

[24] Sa Ribero M, Jouvenet N, Dreux M, Nisole S. Interplay between SARS-CoV-2 and the type I interferon response. PLoS Pathog. 2020 Jul 29;16(7):e1008737.

[25] Monk PD, Marsden RJ, Tear VJ, Brookes J, Batten TN, Mankowski M, Gabbay FJ, Davies DE, Holgate ST, Ho LP, Clark T, Djukanovic R, Wilkinson TMA; Inhaled Interferon Beta COVID-19 Study Group. Safety and efficacy of inhaled nebulised interferon beta-1a (SNG001) for treatment of SARS-CoV-2 infection: a randomised, double-blind, placebo-controlled, phase 2 trial. Lancet Respir Med. 2021 Feb;9(2):196–206.

[26] Cadena C, Ahmad S, Xavier A, Willemsen J, Park S, Park JW, Oh SW, Fujita T, Hou F, Binder M, Hur S. Ubiquitin-Dependent and -Independent Roles of E3 Ligase RIPLET in Innate Immunity. Cell. 2019 May 16;177(5):1187–1200.e16.

[27] Zhang L, Alter HJ, Wang H, Jia S, Wang E, Marincola FM, Shih JW, Wang RY. The modulation of hepatitis C virus 1a replication by PKR is dependent on NF-kB mediated interferon beta response in Huh7.5.1 cells. Virology. 2013 Mar 30;438(1):28–36.

[28] Chen Y, Jiao B, Yao M, Shi X, Zheng Z, Li S, Chen L. ISG12a inhibits HCV replication and potentiates the anti-HCV activity of IFN-α through activation of the Jak/STAT signaling pathway independent of autophagy and apoptosis. Virus Res. 2017 Jan 2;227:231–239.

[29] Adesanya TMA, Campbell CM, Cheng L, Ogbogu PU, Kahwash R. C1 Esterase Inhibition: Targeting Multiple Systems in COVID-19. J Clin Immunol. 2021 May;41(4):729–732.

[30] Smith IA, Knezevic BR, Ammann JU, Rhodes DA, Aw D, Palmer DB, Mather IH, Trowsdale J. BTN1A1, the mammary gland butyrophilin, and BTN2A2 are both inhibitors of T cell activation. J Immunol. 2010 Apr 1;184(7):3514–25.

[31] Pfaender S, Mar KB, Michailidis E, Kratzel A, Boys IN, V’kovski P, Fan W, Kelly JN, Hirt D, Ebert N, Stalder H, Kleine-Weber H, Hoffmann M, Hoffmann HH, Saeed M, Dijkman R, Steinmann E, Wight-Carter M, McDougal MB, Hanners NW, Pöhlmann S, Gallagher T, Todt D, Zimmer G, Rice CM, Schoggins JW, Thiel V. LY6E impairs coronavirus fusion and confers immune control of viral disease. Nat Microbiol. 2020 Nov;5(11):1330–1339.

[32] Zhao X, Zheng S, Chen D, Zheng M, Li X, Li G, Lin H, Chang J, Zeng H, Guo JT. LY6E Restricts Entry of Human Coronaviruses, Including Currently Pandemic SARS-CoV-2. J Virol. 2020 Aug 31;94(18):e00562–20.

