## supplementary Figure S1-3 for "Verification of SARS-CoV-2-Encoded small RNAs and contribution to Infection-Associated lung inflammation"

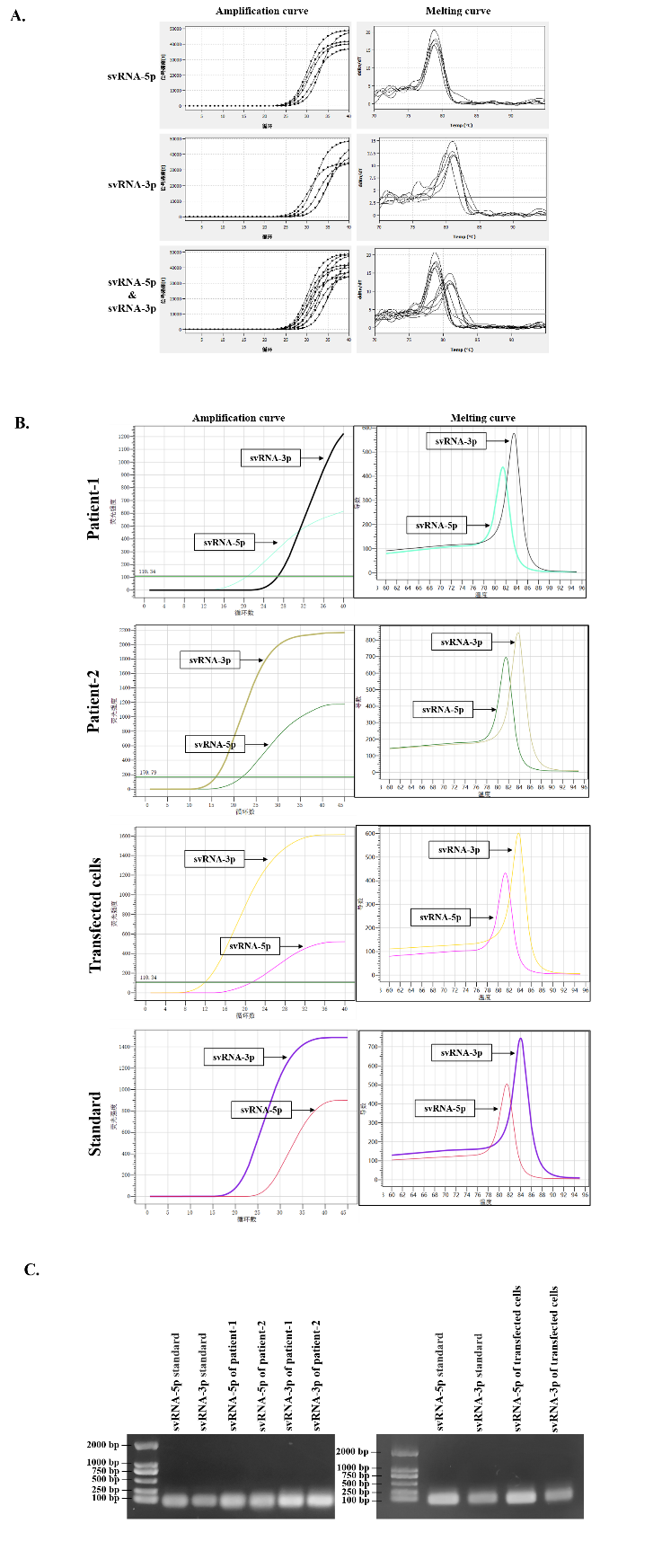

**Figure S1: Verification of the existence of two svRNAs encoded by SARS-CoV-2.**

(A) The amplification and melting curves of the two svRNAs probably maturing from the same precursor named as svRNA-5p and svRNA-3p in 5 nucleic acid samples from pharyngeal swabs of COVID-19 patients obtained from Jiangsu Provincial Center for Disease Control and Prevention.

(B) The corresponding amplification and melting curves for stem-loop RT-PCR performed in FFPE explanted lungs from lung transplantation of two COVID-19 patients, svRNAs precursor transfected 16HBE cells (human lung epithelial cell line) and svRNAs standards, respectively.

(C) The stem-loop RT-PCR products purified in agarose gels before pyrosequencing.

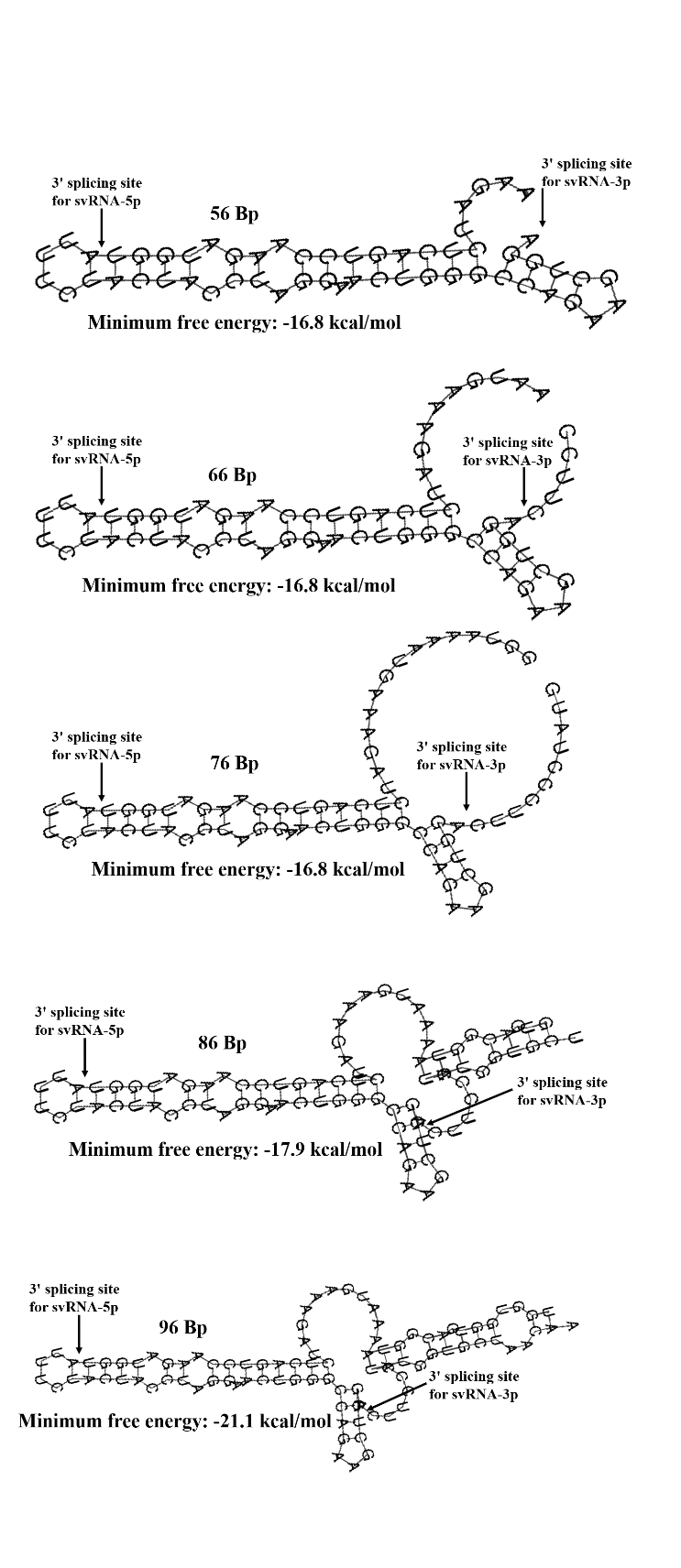

**Figure S2:** The secondary structure of svRNAs precursor with the different lengths (from 56 bp to 96 bp), predicted by RNAfold webserver based on minimum free energy, showed that the splicing sites of the two SARS-CoV-2 encoded svRNAs (svRNA-5p and svRNA-3p) were always stable, no matter with the length of svRNAs precursor.

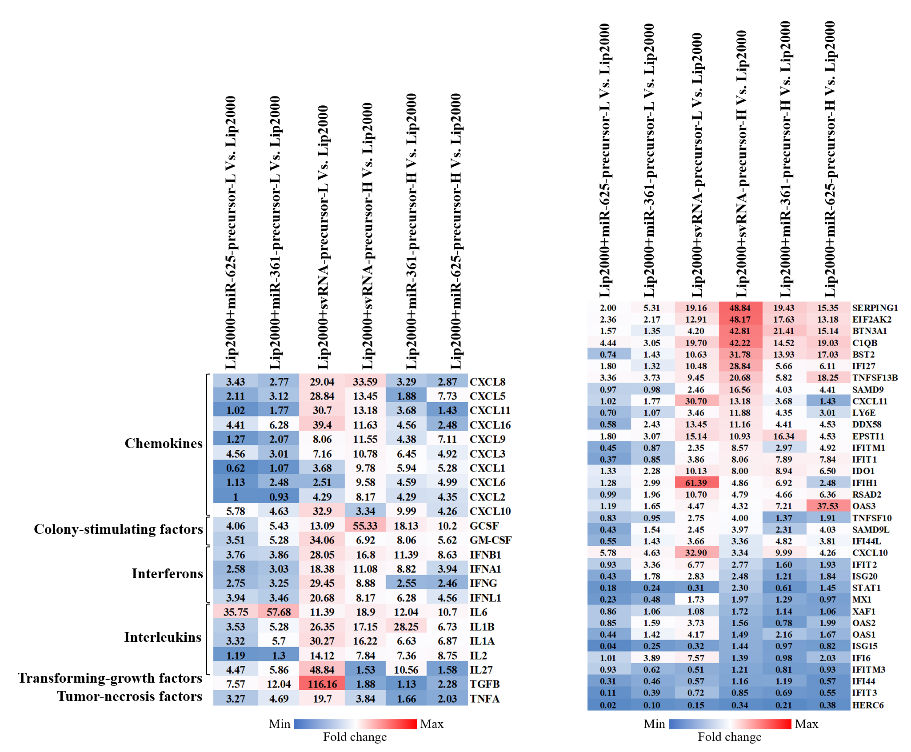

**Figure S3**: 16HBE cells were transfected with control endogenous short RNA precursors and svRNAs precursor at the high (H) and low (L) dosage of 40 pmol and 80 pmol, respectively. The heatmaps showed all the RT-qPCR assay analysis on cells 48 hours post-infection of different dosage of short RNA precursors. The intensity of the color scheme was scaled to relative expression values (fold change). However, no obvious dose-dependence between svRNA precursor and inflammation reaction at transcriptional level was observed.
