## supplementary Table S1-10 for "Verification of SARS-CoV-2-Encoded small RNAs and contribution to Infection-Associated lung inflammation"

**Table S1: Proposal of six potential svRNAs encoded by SARS-CoV-2.**

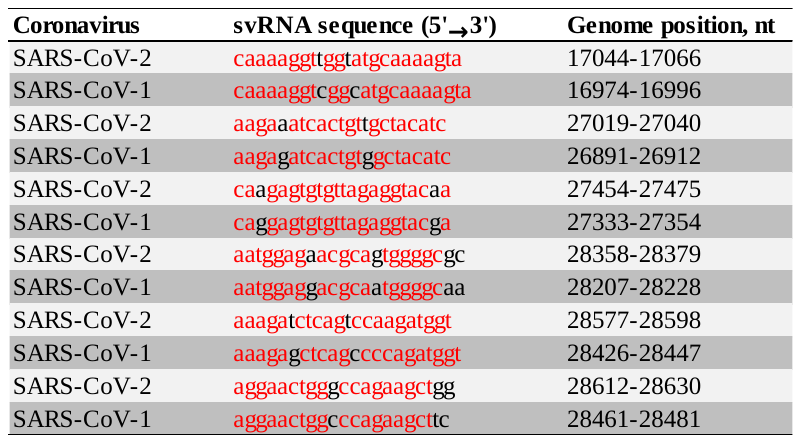

These 6 potential svRNAs derived from the SARS-CoV-2 genome showed significant similarity with those encoded by SARS-CoV-1 in both nucleic acid sequences and virus genome positions, served as candidate SARS-CoV-2-Encoded svRNAs for further verification.

**Table S2: The sequence of forward primers of Poly(A) RT-PCR for the six potential svRNAs encoded by SARS-CoV-2.**

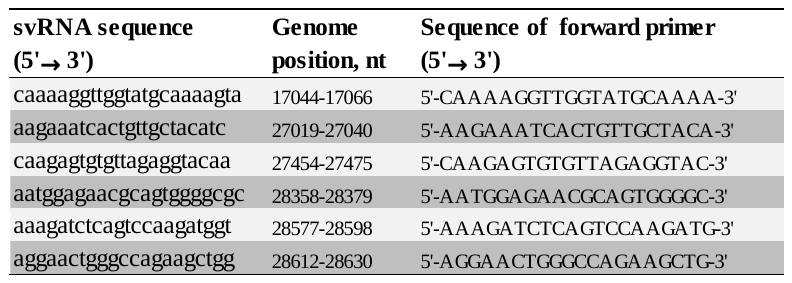

The products of the Poly(A) RT-PCR were then pyrosequenced to verify the existence of the above six potential svRNAs derived from SARS-CoV-2 genome.

**Table S3: The primer sequences of SYBR Green based stem-loop RT-PCR for the two identified svRNAs.**

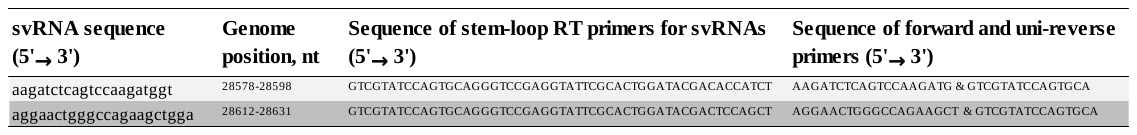

The products of the stem-loop RT-PCR were then pyrosequenced to further confirm the existence of the two identified svRNAs derived from SARS-CoV-2 genome.

**Table S4: The sequences of sense and antisense single-stranded DNA (ssDNA) to form the DNA templates for T7 RNA synthesis.**

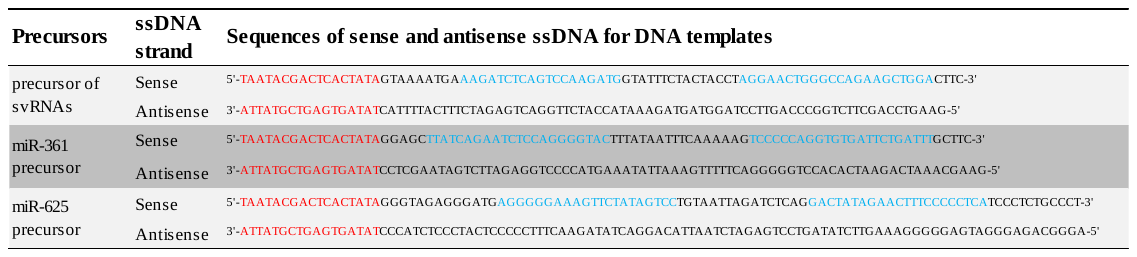

The sequence of T7 promoter region was shown in red. The sequence of the mature short RNAs was shown in blue.

**Table S5: The GEO accession IDs, sample characteristics and experiment types of the 9 datasets associated with COVID-19.**

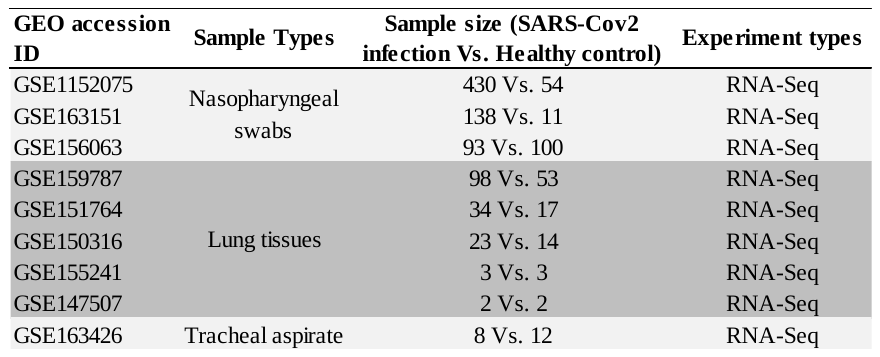

**Table S6: The sequence of primers for 23 cytokine genes covering most of the members of the cytokine storm potentially associated with covid-19.**

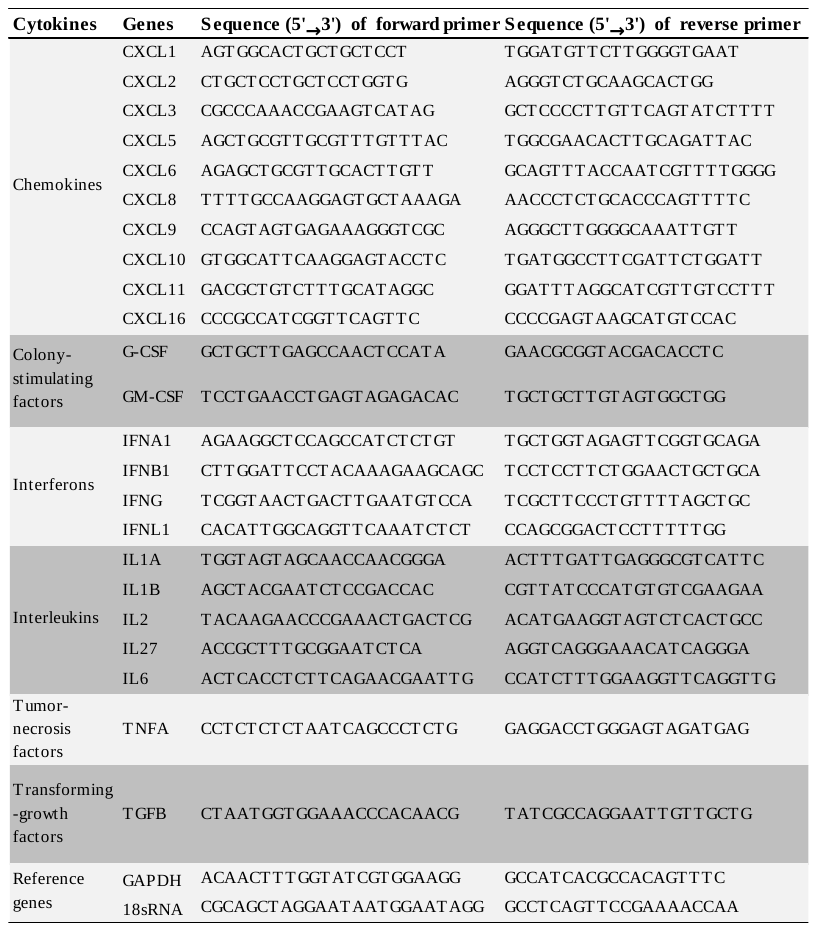

GAPDH and 18S rRNA were considered as reference genes for normalization.

**Table S7: The sequence of primers for genes of the characteristic expression profiling of COVID-19.**

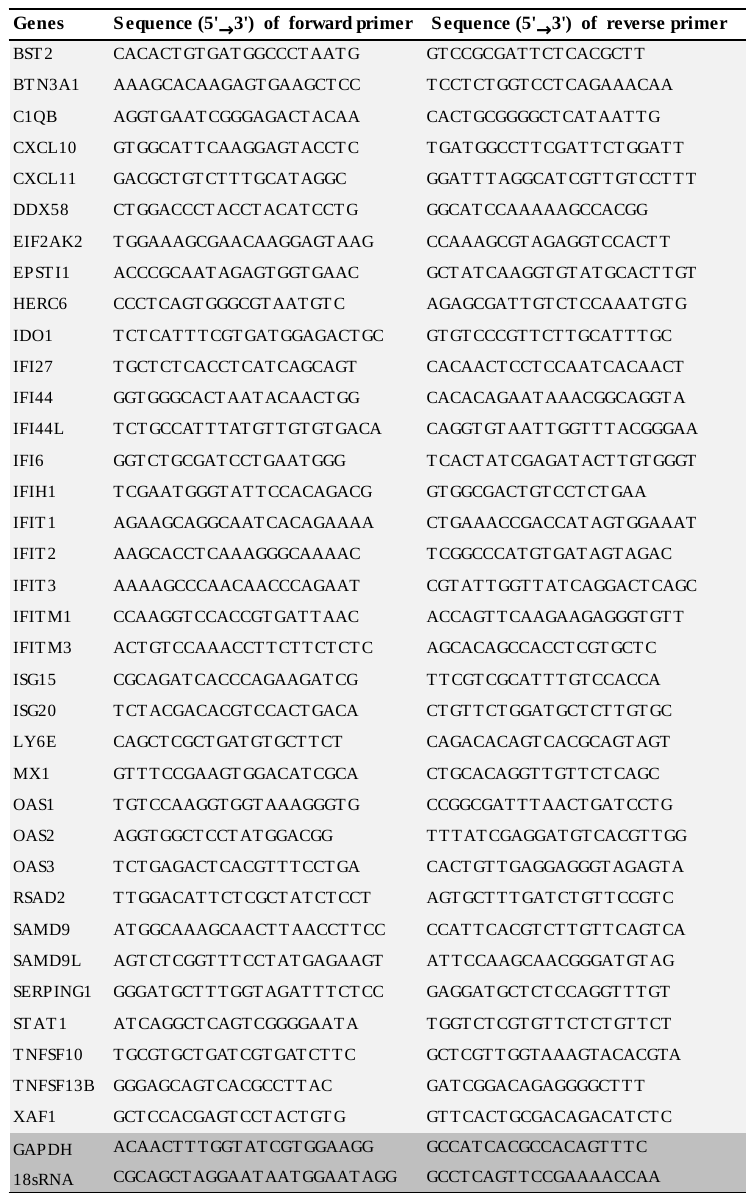

GAPDH and 18S rRNA were considered as reference genes for normalization.

**Table S8: The potential sequence of svRNAs precursor with different length for RNAfold web server to predict the secondary structures.**

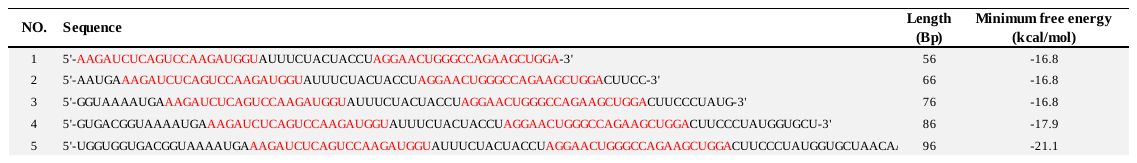

The sequence of the two verified mature svRNAs was shown in red.

**Table S9: The differential expression analysis of the 23 cytokine genes in the lung tissues between the two COVID-19 patients and the controls, respectively.**

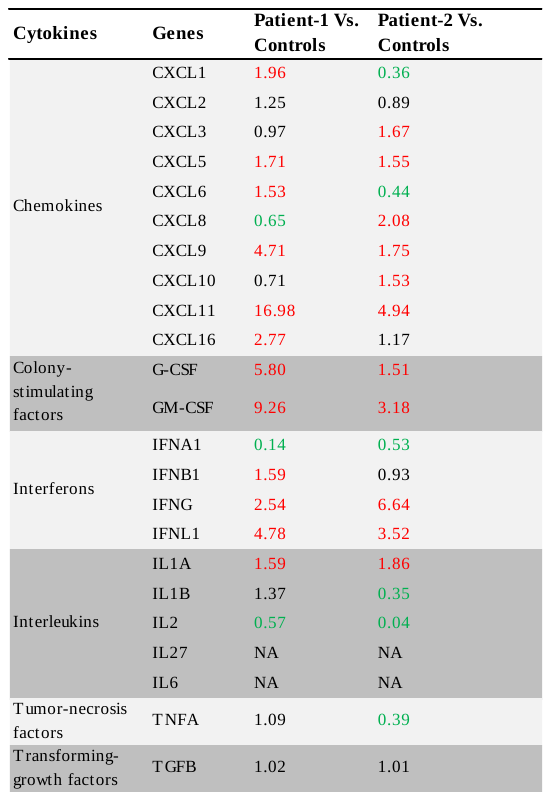

The differential expression level was shown through the average fold changes (FC) between the two COVID-19 patients and the controls. Specifically, FC>1.5 (red) or FC<0.66 (green) were considered as up-regulated or down-regulated in COVID-19, respectively.

**Table S10: The differential analysis of the characteristic expression profiling in the lung tissues between the two COVID-19 patients and the controls, respectively.**

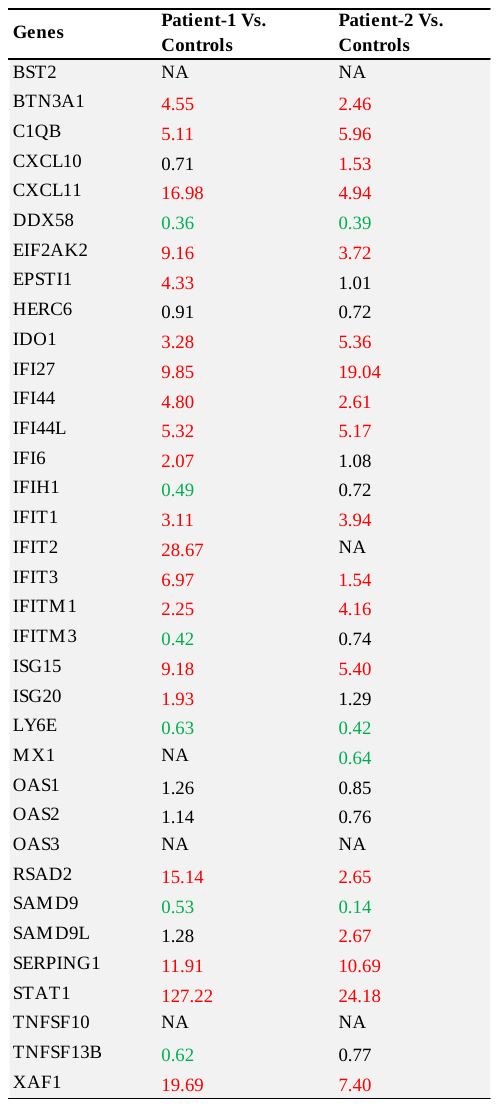

The differential expression level was shown through the average fold changes (FC) between the two COVID-19 patients and the controls. Specifically, FC>1.5 (red) or FC<0.66 (green) were considered as up-regulated or down-regulated in COVID-19, respectively.
